# Durable protection against SARS-CoV-2 Omicron induced by an adjuvanted subunit vaccine

**DOI:** 10.1101/2022.03.18.484950

**Authors:** Prabhu S. Arunachalam, Yupeng Feng, Usama Ashraf, Mengyun Hu, Venkata Viswanadh Edara, Veronika I. Zarnitsyna, Pyone Pyone Aye, Nadia Golden, Kristyn W. M. Green, Breanna M. Threeton, Nicholas J. Maness, Brandon J. Beddingfield, Rudolf P. Bohm, Jason Dufour, Kasi Russell-Lodrigue, Marcos C. Miranda, Alexandra C. Walls, Kenneth Rogers, Lisa Shirreff, Douglas E Ferrell, Nihar R. Deb Adhikary, Jane Fontenot, Alba Grifoni, Alessandro Sette, Derek T. O’Hagan, Robbert Van Der Most, Rino Rappuoli, Francois Villinger, Harry Kleanthous, Jay Rappaport, Mehul S. Suthar, David Veesler, Taia T. Wang, Neil P. King, Bali Pulendran

## Abstract

Despite the remarkable efficacy of COVID-19 vaccines, waning immunity, and the emergence of SARS-CoV-2 variants such as Omicron represents a major global health challenge. Here we present data from a study in non-human primates demonstrating durable protection against the Omicron BA.1 variant induced by a subunit SARS-CoV-2 vaccine, consisting of RBD (receptor binding domain) on the I53-50 nanoparticle, adjuvanted with AS03, currently in Phase 3 clinical trial (NCT05007951). Vaccination induced robust neutralizing antibody (nAb) titers that were maintained at high levels for at least one year after two doses (Pseudovirus nAb GMT: 2207, Live-virus nAb GMT: 1964) against the ancestral strain, but not against Omicron. However, a booster dose at 6-12 months with RBD-Wu or RBD-β (RBD from the Beta variant) displayed on I53-50 elicited equivalent and remarkably high neutralizing titers against the ancestral as well as the Omicron variant. Furthermore, there were substantial and persistent memory T and B cell responses reactive to Beta and Omicron variants. Importantly, vaccination resulted in protection against Omicron infection in the lung (no detectable virus in any animal) and profound suppression of viral burden in the nares (median peak viral load of 7567 as opposed to 1.3×10^7^ copies in unvaccinated animals) at 6 weeks post final booster. Even at 6 months post vaccination, there was significant protection in the lung (with 7 out of 11 animals showing no viral load, 3 out of 11 animals showing ~20-fold lower viral load than unvaccinated controls) and rapid control of virus in the nares. These results highlight the durable cross-protective immunity elicited by the AS03-adjuvanted RBD-I53-50 nanoparticle vaccine platform.

## Introduction

Waning immunity coupled with the continuing emergence of immune evasive variants represents a significant challenge in managing the COVID-19 pandemic. The efficacy of even the most effective mRNA vaccines decreased 20 – 30% by six months post-two-dose vaccine series (Feikin et al., 2022; Goldberg et al., 2021). The efficacy declined more precipitously against Omicron, a variant highly resistant to vaccine-induced therapeutic antibodies (Cameroni et al., 2022; Dejnirattisai et al., 2022; Hoffmann et al., 2022), reaching inadequate (8.8% following 2-dose Pfizer-BioNTech mRNA vaccination) to no protection (ChAdOx1-nCOV19) by 5 – 6 months following vaccination (Andrews et al., 2022). The waning efficacy thus mandates a booster vaccination while ~40% of the world’s population is yet to receive full vaccination, resulting in a large gap in vaccine equity.

We recently reported a study in which we compared the immunogenicity and protective efficacy of the RBD-I53-50 nanoparticle immunogen formulated with five different adjuvants in NHPs (Arunachalam et al., 2021). AS03, an oil-in-water emulsion containing α-tocopherol, adjuvanted RBD-I53-50 vaccination elicited the most potent and broad nAb response, as well as substantial T cell responses, and conferred significant protection against SARS-CoV-2 challenge in the upper and lower airways. More importantly, the vaccine-induced nAbs persisted for at least 6 months indicating durability of immune responses induced by the adjuvanted subunit vaccine platform. Here, we evaluated the durability of immune protection after a final booster immunization with RBD-Wu or RBD-β against the immune-evasive Omicron variant.

## Results

### Study design

The study involved four groups of male Rhesus macaques. The first group of 5 animals were immunized thrice with RBD-Wu + AS03, at days 0 and 21 followed by a final booster ~6 months later, mimicking the human population that received three doses of mRNA vaccines (**Fig. 1a**, group RBD-Wu/RBD-Wu/RBD-Wu). The second and third groups were from our previous study (Arunachalam et al., 2021) in which one group of 5 animals received two doses of RBD-I53-50-Wu (RBD-Wu or RBD-NP-Wu) and the other group comprising 6 animals received two doses of HexaPro-I53-50 (HexaPro or HexaPro-NP-Wu). Both immunogens were administered with the AS03 adjuvant on days 0 and 21 using a prime-boost regimen. In the current phase of the study, all 11 animals from both groups were boosted with an I53-50 nanoparticle immunogen displaying RBD-β stabilized with the Rpk9 mutations (Ellis et al., 2021) approximately a year after the first immunization series (**Fig. 1a**, groups RBD-Wu/RBD-Wu/RBD-β and HexaPro/HexaPro/RBD-β). The Beta variant was selected on the grounds that it was a prevalent and one of the most antibody-evading SARS-CoV-2 strains at the time of the study design. Therefore, we had planned to challenge these animals with the Beta variant originally. However, the emergence of Omicron as the dominant strain while the study was ongoing prompted us to challenge the animals with Omicron instead, and assess heterologous protection. The fourth group of 5 animals were unvaccinated controls. To assess immunogenicity, the RBD-Wu/RBD-Wu/RBD-Wu animals were followed longitudinally from the day of the first immunization whereas the two groups from the previous study were followed from the day of the final booster (**Fig. 1a**). To assess protective efficacy, we challenged the animals with 2×10^6^ plaque-forming units of SARS-CoV-2 Omicron virus. Notably, we challenged the first group at ~6 weeks after the final booster and the other two groups at ~6 months after the final booster to assess protection at the peak of immune responses versus when immune responses were waning.

**Figure 1.**
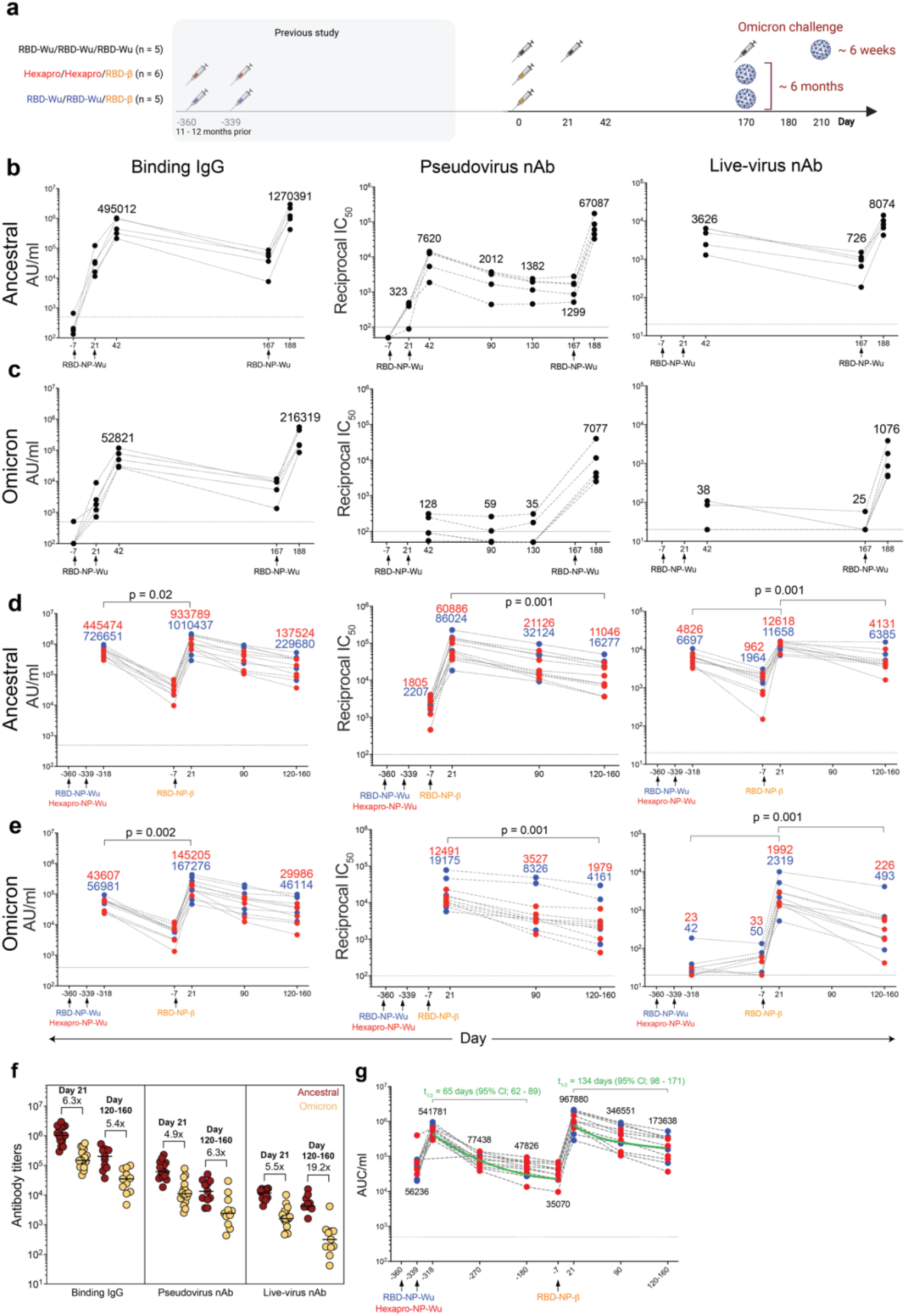
Serum antibody responses induced by AS03-adjuvanted RBD-nanoparticle vaccination. **a**, A schematic of the study design. **b – c**, Anti-Spike binding IgG, pseudovirus nAb titers, and live-virus nAb titers against the ancestral WA1 (**b**), and Omicron (**c**) strains in the RBD-Wu/RBD-Wu/RBD-Wu group (n = 5). **d – e**, Anti-Spike binding IgG, pseudovirus nAb titers, and live-virus nAb titers against the ancestral WA1 (**d**), and Omicron (**e**) strains in the RBD-Wu/RBD-Wu/RBD-β (blue, n = 5) and HexaPro/HexaPro/RBD-β (red, n = 6) groups. The numbers within the graphs show geometric mean titers. **f**, Antibody titers against the ancestral and Omicron strains at the time points (n = 16 on day 21 and 11 on day 120-160) indicated on top. The numbers within the graph indicate fold-change between ancestral and Omicron antibody titers. **g**, Binding antibody titers against the ancestral strain in sera collected at all indicated time points (n = 11). The green lines show the fit using the power-law model to calculate decay rates. The t_1/2_ value after the second dose was calculated until day 180, i.e., 159 days after the second dose. The data in **f** and **g** contain portion of the data contained within **b** to **e**. In all figures, each circle represents an animal. The statistical differences between time points were determined using Wilcoxon matched-pairs signed rank test. For **d** and **e**, the statistical analysis was performed with all 11 animals.

### Humoral immune responses

Vaccination with RBD-Wu + AS03 elicited binding IgG titers against the Spike protein of the ancestral strain, Omicron and Beta variants detectable on day 21. The titers increased greater than 10-fold after the second immunization (GMT: 495012 and 52821 AU/ml against ancestral and Omicron, respectively) (**Fig. 1b, c – left panel and Supplementary Fig. 1a**) and reduced to pre-booster levels by 6 months. The final booster immunization increased the titers by 2.5- and 4-fold against the ancestral (GMT 1270391) and Omicron (GMT 216319) strains, respectively, relative to the titers at the peak of the second immunization (**Fig. 1b, c**). We also observed detectable nAb response against the ancestral strain after one immunization, which increased ~20-fold (GMT 7620) after the second immunization and was maintained at substantial levels (GMT 1299) until the booster dose ~6 months later (**Fig. 1b – middle panel**). In contrast, there was only low levels of nAb response against the Omicron variant (**Fig. 1c, middle panel**). The final booster immunization at ~6 months increased the titers strikingly against the ancestral strain (GMT 67087) as well as Omicron (GMT 7077) and Beta variants (**Fig. 1b, c – middle panel and Supplementary Fig. 1b – left panel**). Consistent with pseudovirus nAb response, vaccination also induced live-virus nAb titers against the ancestral (GMT 3626) and Beta (GMT 377) strains but not Omicron after two immunizations, comparable to the responses seen in our previous study(Arunachalam et al., 2021) (**Fig. 1b, c – right panel and Supplementary Fig. 1b – right panel**). The titers against ancestral strain decreased ~5-fold by 6 months (GMT 726 against the ancestral strain) when the booster was administered. The booster vaccination enhanced the responses against all three strains reaching peak geometric mean titers of 8074, 1076 and 4527 against the ancestral strain, Omicron and Beta variants, respectively (**Fig. 1b, c – right panel and Supplementary Fig. 1b – right panel**).

In the two groups that were boosted with RBD-β at approximately a year after the two-dose primary vaccine series (**Fig. 1a**), the booster vaccination elicited binding IgG responses against ancestral, Omicron and Beta strains comparable to the titers in the RBD-Wu/RBD-Wu/RBD-Wu group. The responses were maintained durably through the ~5-month follow-up period (GMT 967880 and 173638 at peak and ~5-month time points, respectively, against the ancestral strain) (**Fig. 1d, e – left panel and Supplementary Fig. 1c)**. Since we observed no significant difference in antibody titers between the RBD-I53-50 and HexaPro-I53-50 groups prior to (as in our previous study (Arunachalam et al., 2021)) or after the booster immunization, the GMTs presented here represent the geometric mean of all 11 animals combined from both groups. Consistent with binding IgG titers, neutralization activity against the ancestral pseudovirus (GMT 1978) and live-virus (GMT 1331) was still detectable prior to the final booster at ~1 year after the two-dose primary vaccine series (**Fig. 1d, middle and right panels**). The RBD-β booster immunization enhanced the titers significantly against the ancestral strain, Omicron and Beta variants. The geometric mean titers against the ancestral strain reached as high as 71244 and 12172 (in the 11 animals combined from the RBD-Wu/RBD-Wu and HexaPro/HexaPro groups) in the pseudovirus and live-virus neutralization assays, respectively. Boosting with RBD-Wu or RBD-β elicited comparable titers in both groups against each of the three viral strains measured (**Supplementary Fig. 1e**). While the pseudovirus nAb titers decreased only by ~5-fold against all three strains, the live-virus nAb titers decreased by ~2.5 and ~7-fold against the ancestral and Omicron strains, respectively, over the ~5-month follow-up period (**Fig. 1d, e middle and right panels and Supplementary Fig. 1d)**. In addition, we also assessed binding and live-virus nAb titers in sera collected on day 21 post second vaccination (represented as day −318 in the figures, relative to the final booster dose) to determine the durability of the antibody response after the second dose to compare the durability and magnitude of the antibody responses after second and third doses. While there was only low levels of nAb response against Omicron after the second dose (consistent with the responses in the RBD-Wu/RBD-Wu/RBD-Wu group), the live-virus nAb titers against the ancestral strain reduced only less than 10-fold within the 1 year before the final booster vaccination (**Fig. 1d, right panel**) (**Fig. 1d, right panel**). Finally, we examined the reduction in nAb titers against Omicron in comparison with the ancestral strain. Consistent with several recent studies (Cameroni et al., 2022; Cheng et al., 2022; Edara et al., 2022; Garcia-Beltran et al., 2022; Pajon et al., 2022; Schmidt et al., 2022; Sievers et al., 2022; Walls et al., 2022), the antibody titers were ~5-fold lower to the Omicron variant than was elicited against the ancestral strain (**Fig. 1f**). Although the magnitude of nAbs measured using the pseudovirus assay was 10-fold higher than the live-virus neutralization assays, the titers correlated with each other and with binding antibodies significantly (**Supplementary Fig. 1f**).

Next, we estimated the half-life of binding IgG antibodies using the exponential decay and power law decay models. The exponential decay model assumes a constant decay rate over time, and the power law decay model assumes that decay rates decrease over time. The power law model fitted significantly better compared to the exponential decay model after the second vaccine dose (ΔBICc = 24.2), and there was no statistically significant difference between the fit of the exponential and power-law models after the third dose (ΔBICc = 2.5). Therefore, we compared the half-lives after second and third doses calculated using the power-law decay model, which showed that the half-life after the third dose was ~2-fold longer than after the second dose. The estimated half-lives against the ancestral strain after the second and third doses were 65 days (95% Cl; 62 – 89) and 134 days (95% Cl; 98 – 171), respectively (**Fig. 1g**). Of note, the t_1/2_ values presented in **Fig. 1g** were calculated using data from equivalent time points after the 2^nd^ and 3^rd^ doses (see methods for details). Collectively, these data demonstrate that the RBD-I53-50 immunogen adjuvanted with AS03 stimulates robust and durable nAb response against multiple SARS-CoV-2 variants including Omicron. In addition, there was no significant difference in the antibody responses elicited by RBD-Wu or RBD-β booster immunizations, consistent with recent studies assessing variant booster vaccination (Choi et al., 2021; Corbett et al., 2021).

### Mucosal antibody responses

Antibody responses at the mucosa are critical to developing protective immunity against SARS-CoV-2. To assess if this vaccine regimen elicits antibody responses at the mucosa, we measured binding antibody titers in the bronchoalveolar lavage (BAL) collected longitudinally from vaccinated animals using the mesoscale platform. Consistent with the serum antibody responses, RBD-Wu + AS03 immunization resulted in significant induction of antibodies that bound to the ancestral (GMT: 108 AU/μg total IgG) as well as Omicron (GMT: 15 AU/μg) Spike proteins after two doses (**Fig. 2a**). The final dose boosted the titers by ~2-fold and ~4-fold against ancestral (GMT 233 AU/μg) and Omicron (56 AU/μg) variants, respectively. Similarly, the animals in the RBD-Wu/RBD-Wu or HexaPro/HexaPro that were boosted with RBD-β elicited titers comparable to that of the RBD-Wu/RBD-Wu/RBD-Wu group and maintained durably up to 4 – 5 months after vaccination with only a ~4-fold reduction in titers (**Fig. 2a**). There was also a significant correlation between binding antibody titers in serum and BAL fluid (**Fig. 2b**).

**Figure 2.**
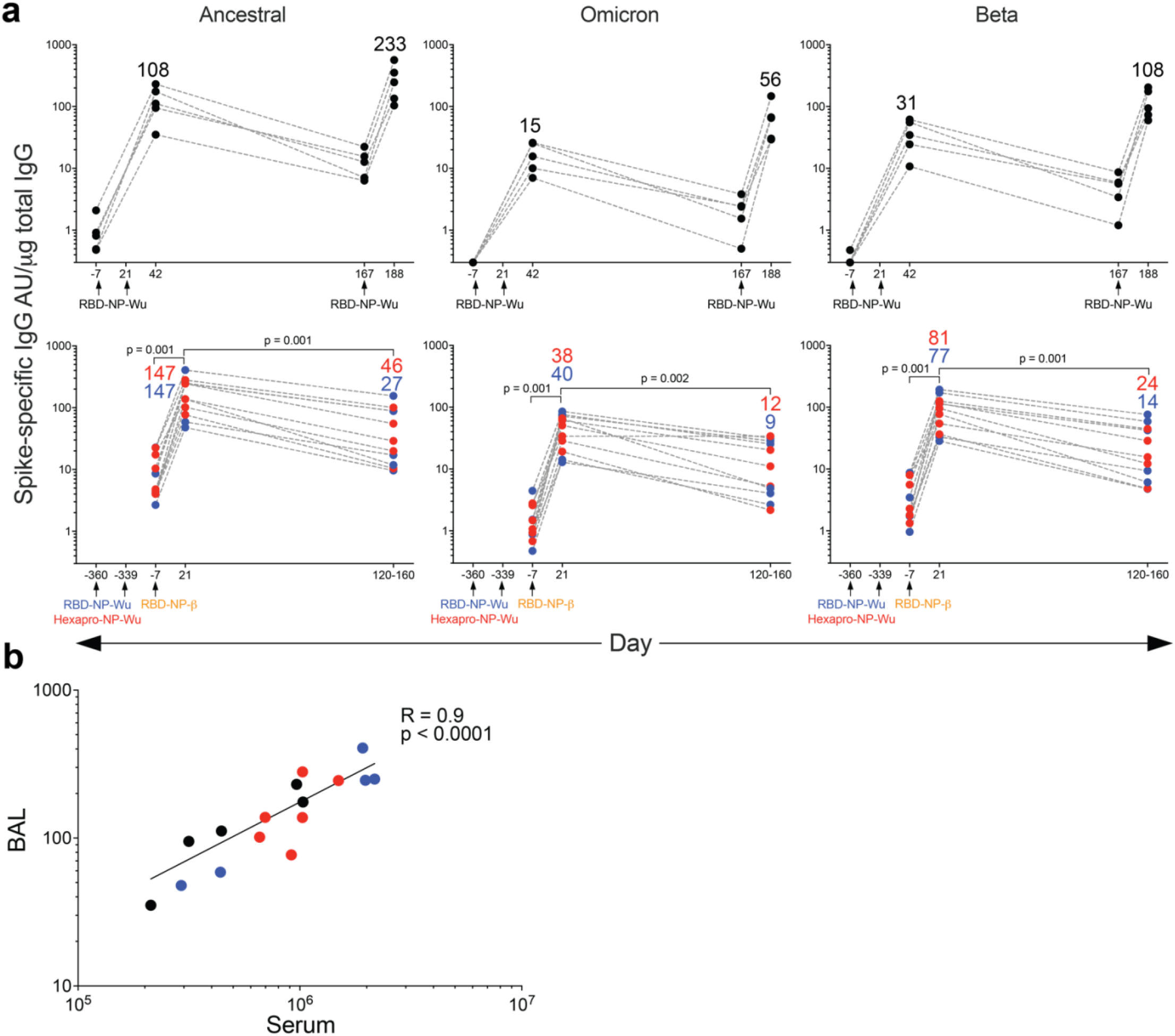
Mucosal antibody responses induced by AS03-adjuvanted RBD-nanoparticle vaccination. **a**, Spike-binding IgG titers against the ancestral strain (left panel), Omicron (middle panel) and Beta (right panel) variants in the BAL fluid of animals in the RBD-Wu/RBD-Wu/RBD-Wu (top panels, n = 5) and RBD-Wu/RBD-Wu/RBD-β (blue, n = 5) and HexaPro/HexaPro/RBD-β (red, n = 6) groups (bottom panels). The statistical differences between time points were determined using Wilcoxon matched-pairs signed rank test combining 11 animals from the RBD-Wu/RBD-Wu/RBD-β and HexaPro/HexaPro/RBD-β groups. **b**, Spearman’s correlation between binding IgG measured at 21 days post final booster in serum versus BAL fluid.

### Cellular immune responses

We measured T cell responses using intracellular cytokine staining (ICS) assay following a 6 h stimulation of peripheral blood mononuclear cells (PBMCs) with overlapping peptide pools spanning the Spike proteins of the ancestral, Omicron and Beta variants (Tarke et al., 2022). Consistent with our previous study(Arunachalam et al., 2021), RBD-Wu + AS03 vaccination elicited CD4 T cell responses, both Th1 and Th2 type, after two doses (**Fig. 3a, b**). The responses subsided to near baseline levels by 6 months, which was boosted by the final booster immunization (**Fig. 3a, b and Supplementary Fig. 2a**). Similarly, the final booster immunization with RBD-β in the other two groups also increased antigen-specific CD4 T cell responses (Th1 as well as Th2) (**Fig. 3c, d and Supplementary Fig. 2b**). The responses decreased significantly by 4 - 5 months post final booster immunization but were still detectable. The polyfunctional profile of Spike-specific CD4 T cells after two doses was comparable to our previous study(Arunachalam et al., 2021), with the majority (~70%) of the cells expressing IL-2 +/− TNF, and a balanced Th1/Th2 profile expressing IFN-γ or IL-4 (**Fig. 3e**). The profile after the third dose was similar; however, the cells expressing IL-2 and/or TNF without IFN-γ or IL-4 were reduced marginally (40% - 60% in contrast to 70%) (**Fig. 3e**). Of note, the CD4 T cell response to Omicron was reduced only marginally (15%), albeit significantly, in comparison to responses to the ancestral strain (**Fig. 3a, b, f**), consistent with studies in humans (Gao et al., 2022; Keeton et al., 2022; Tarke et al., 2022). In addition to Th1- and Th2-cytokines, AS03 also promotes follicular T helper (T_FH_) response. We found low but detectable IL-21 and high levels of CD154 in response to vaccination (**Fig. 3g**). In summary, the booster immunization with RBD-Wu or RBD-β elicited considerable CD4 T cell responses with only a marginal reduction in Omicron- or Beta-specific T cell responses.

**Figure 3.**
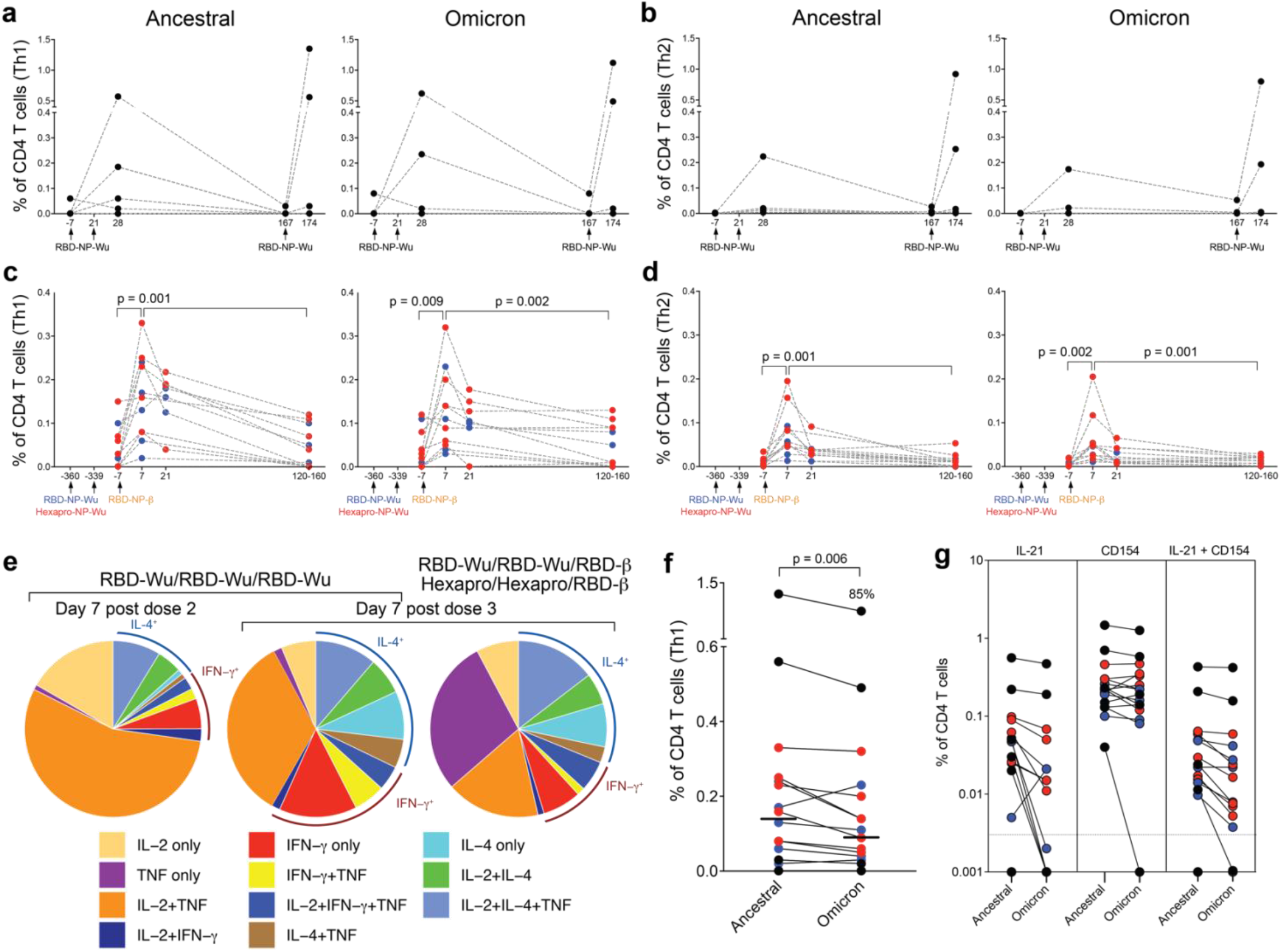
T cell responses induced by AS03-adjuvanted RBD-nanoparticle vaccination. **a - b**, Frequency of Spike-specific CD4 T cell responses against ancestral (left panel) and Omicron (right panel) strains in the RBD-Wu/RBD-Wu/RBD-Wu group. **c – d**, Frequency of Spike-specific CD4 T cell responses against ancestral (left panel) and Omicron (right panel) strains in the RBD-Wu/RBD-Wu/RBD-β (blue) and HexaPro/HexaPro/RBD-β (red) groups. CD4 T cells secreting IL-2, IFN-γ, or TNF are plotted as Th1-type responses (**a, c**) and IL-4-producing CD4 T cells are shown as Th2-type responses (**b, d**) The statistical differences between time points were determined using Wilcoxon matched-pairs signed rank test. **e**, Pie charts representing the proportions of RBD-specific CD4 T cells expressing one, two, or three cytokines as shown in the legend. **f**, Comparison of CD4 T cell frequencies between ancestral and Omicron viral strains measured on day 7 post final booster immunization. The statistical difference was determined using Wilcoxon matched-pairs signed rank test. The % value on top of Omicron represents the proportion of Omicron-specific responses relative to responses against the ancestral strain. **g**, Spike-specific IL-21^+^, CD154^+^, and CD154^+^IL-21^+^ CD4 T cell responses measured in blood on day 7 post final booster immunization. In all plots, each circle represents an animal, In **f** and **g**, black, blue, and red colors indicate RBD-Wu/RBD-Wu/RBD-Wu, RBD-Wu/RBD-Wu/RBD-β and HexaPro/HexaPro/RBD-β groups, respectively.

Next, we assessed Spike-specific memory B cells by flow cytometry analysis of PBMCs labelled with fluorescent-tagged Spike protein of the ancestral, Omicron and Beta variants (**Fig. 4a and Supplementary Fig. 3a**). Immunization with RBD-Wu + AS03 elicited robust Spike-specific memory B cells (up to 1%) 21 days after the second immunization, more than 50% of which bound to all three probes (**Fig. 4b, c**). The memory B cells consisted of 0.021% and 0.011% of all CD20^+^ B cells 6 months and 1 year after two doses, respectively (**Fig. 4b**). The third dose, either with RBD-Wu or RBD-β boosted the memory B cell frequencies by ~10-fold with a phenotype transition from resting (CD21^+^/CD27^+^) to activated (CD21^−^/CD27^+^) memory (**Fig. 4a, b and Supplementary Fig. 3b**) and an increased proportion of memory B cells binding to all three probes (**Fig. 4c**). The frequency of total Spike-specific memory B cells decreased gradually and reached levels close to the pre-booster time point by 4 - 5 months (**Fig. 4b**). Together, these data show that two doses of immunization elicited a durable memory B cell response and the booster immunization with RBD-Wu or RBD-β further broaden the cross-reactivity of memory B cells to SARS-CoV-2.

**Figure 4.**
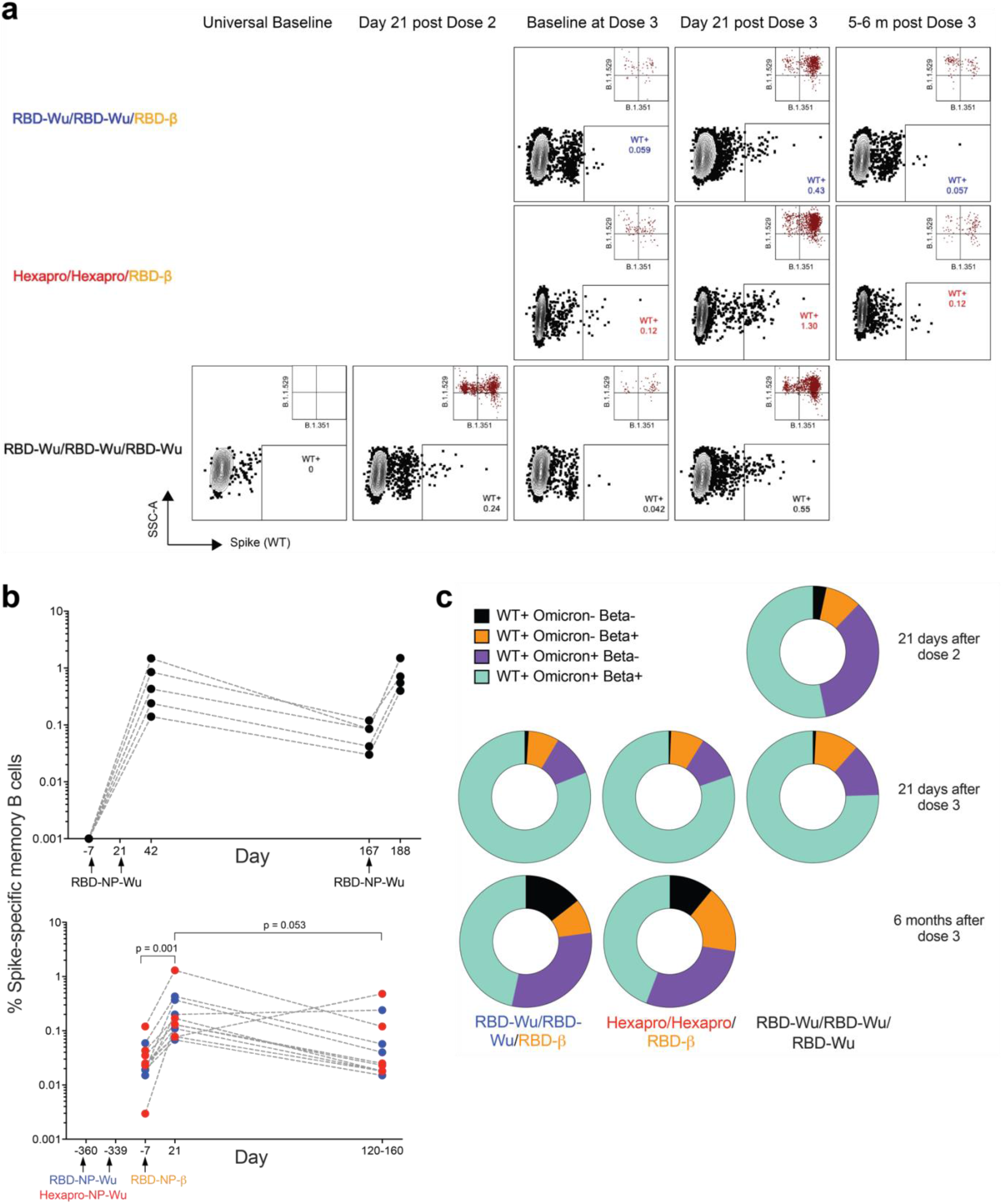
Memory B cell responses induced by AS03-adjuvanted RBD-nanoparticle vaccination. **a**, A representative flow cytometry profile showing ancestral Spike-specific B cell frequencies gated as live, CD20^+^ IgD^−^ IgM^−^ Spike^+^ cells. The insets show proportion of Wu Spike^+^ cells that bind to Omicron versus Beta probes. A higher number of events is shown in the insets to improve visibility. **b**, Frequency of Spike-specific memory B cells relative to CD20^+^ B cells in the RBD-Wu/RBD-Wu/RBD-Wu (top panel), RBD-Wu/RBD-Wu/RBD-β (blue, bottom panel) and HexaPro/HexaPro/RBD-β groups (red, bottom panel). **c**, Donut charts showing proportion of Spike-specific memory B cells bound to ancestral (WT), Omicron and Beta probes as indicated in the legend.

### Protection against Omicron

Omicron is characterized by high resistance to vaccine-induced nAbs and therapeutic monoclonal nAbs (Cameroni et al., 2022; McCallum et al., 2022), which results in a striking decline in vaccine efficacy against Omicron (Andrews et al., 2022). Booster vaccination has been recommended to improve effectiveness (Omer and Malani, 2022); however, the efficacy against symptomatic infection remains at ~65% at 2 – 4 weeks after a booster vaccination with BNT162b2 or mRNA1273 (Accorsi et al., 2022; Andrews et al., 2022). These findings prompted us to examine protection at the peak time point in our study in addition to evaluating protection at 6 months when the vaccine-induced immunity is waning. Therefore, we challenged the animals receiving RBD-Wu/RBD-Wu/RBD-Wu + AS03 at 6 weeks post final booster. Given that the immune responses in the animals belonging to the other two groups (RBD-Wu/RBD-Wu/RBD-β and HexaPro/HexaPro/RBD-β) were comparable, we challenged all 11 animals at 6 months post final booster (**Fig. 1a**). All the animals were challenged with 2×10^6^ PFU via the intranasal (1×10^6^ PFU) and intratracheal (1×10^6^ PFU) routes.

Two days after challenge, 4 out of 5 unvaccinated animals had a subgenomic viral load of 28,000 – 150,000 copies/ml (N gene) in the BAL fluid. The viral load persisted until day 7 and reduced to undetectable levels by day 14. In contrast, the animals challenged at the peak of immune responses (RBD-Wu/RBD-Wu/RBD-Wu group) demonstrated complete protection in the lung, with none of the five animals showing a detectable viral load at any time point in the BAL fluid (**Fig. 5a**). Of the 11 animals challenged at ~6 months post final booster vaccination, there was incomplete but significant protection in the lung compartment. Seven of the eleven animals were completely protected while the remaining 4 animals (2 animals from each vaccination group further supporting equivalent immunogenicity in these groups) had a peak viral load of 1135, 3114, 4195 and 33845 copies/ml, as opposed to a median viral load of ~59,000 copies/ml in the unvaccinated controls. Of the 4 animals showing a viral load in BAL, there was rapid viral control with only one animal showing a viral load by day 7 (**Fig. 5a**).

**Figure 5.**
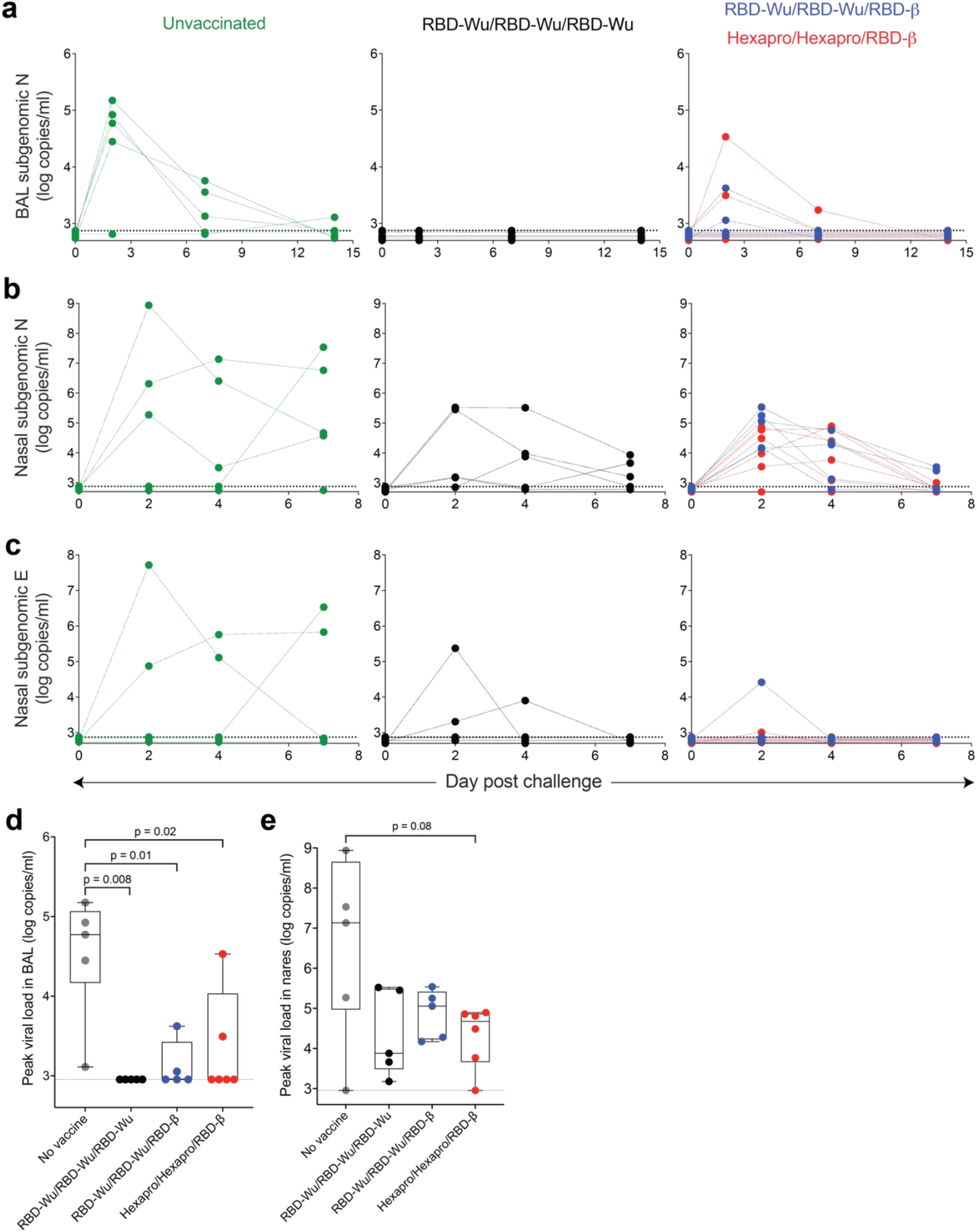
Protection against SARS-CoV-2 Omicron challenge. **a**, SARS-CoV-2 Omicron viral load in BAL fluid measured using N gene subgenomic PCR. **b – c**, SARS-CoV-2 Omicron viral load in nasal swabs measured using N (**b**) and E (**c**) gene subgenomic PCRs. **d - e**, Peak N gene viral load in the BAL fluid (**d**) and nasal swabs (**e**). The statistical differences between groups were determined using Mann-Whitney test.

To assess protection in the upper airways, we assessed viral loads in nasal swabs using N and E gene subgenomic PCRs. Four of the five control animals were infected. The median peak viral load was 1.36×10^7^ copies/ml of the N gene, which was maintained up to day 7 in all 4 infected animals (**Fig. 5b, c, left panel**). In contrast, although all vaccinated animals showed detectable viral loads irrespective of whether they were challenged at peak or 6 months later, the viral burden was rapidly suppressed. The peak median viral loads were 7567 and 64366 copies/ml in the animals challenged at peak and 6 months, respectively, and viral loads were reduced to the levels of detection limit by day 7 (**Fig. 5b, c**). Comparing peak viral loads revealed a statistically significant difference in the BAL fluid between each of the three vaccinated groups compared to unvaccinated controls, and viral loads in the nasal compartment were substantially reduced (**Fig. 5d, e**). Collectively, these data demonstrate that the RBD-I53-50 nanoparticle vaccine adjuvanted with AS03 confers impressive protection against Omicron even at 6 months after the final booster immunization.

## Discussion

Waning immunity, especially against Omicron, represents an enormous challenge in managing the ongoing pandemic. The efficacy against symptomatic infection of the Pfizer-BioNTech mRNA vaccine, one of the most efficacious vaccines against SARS-CoV-2, declined from >90% to 67% against Omicron 2 – 4 weeks after the second booster vaccination. The efficacy further declined to 45% in the following 10 weeks demonstrating the effect of rapidly waning immunity (Andrews et al., 2022). We show here that the RBD-I53-50 + AS03 adjuvanted nanoparticle vaccine confers complete protection against Omicron infection in the lung at 6 weeks post final booster and ~65% protection at 6 months post final booster. Recent studies in NHPs vaccinated with the Moderna mRNA platform (Gagne et al., 2022), or a homologous or a heterologous prime-boost regimen with the Ad26 and BNT162b2 vaccine platforms (Chandrashekar et al., 2022) evaluated protection only at the peak (~1 month) of the vaccine-induced immunity. We also observed protection in the lung and rapid control of virus in the nares; however, our study provides data demonstrating protection even at 6 months post final booster. Whether similar protection is observed in humans remains to be seen.

A second important finding of our study is the remarkable durability of binding and nAb titers elicited by this vaccine platform. Comparison of live-virus nAb titers measured using the same assay by the same laboratory (Edara et al., 2022; Edara et al., 2021) shows that the peak response after two doses of mRNA1273 (live-virus nAb titer GMT: 5560 (Gagne et al., 2022)) and the RBD-I53-50 + AS03 (GMT: 6697) vaccines were comparable; however, the live-neut GMTs were 330 (Gagne et al., 2022) and 1964 at the time of the final booster at ~10 – 11 months post second dose of mRNA1273 and RBD-I53-50+AS03, respectively, suggesting that the nAb titers were relatively more durable following vaccination with this adjuvanted nanoparticle platform. However, it should be noted that the assays were not performed simultaneously to allow for an ideal head-to-head comparison.

In addition to eliciting durable humoral immune responses, the RBD-I53-50 + AS03 vaccination results in substantial memory T and B cell responses. Vaccination stimulates substantial CD4 T cell responses but little CD8 T cell responses. Despite the lack of CD8 T cell immunity, the protection observed in these animals was comparable to vaccine modalities that elicit CD8 T cells such as the Ad26 platform (Chandrashekar et al., 2022). Furthermore, although there were only low levels of nAb response to Omicron after second immunization, the booster vaccination induced high levels of Omicron-targeting nAbs suggesting that the memory B cells continue to mature in the 1-year interval between second and third vaccination. Whether memory B cells continue to mature after the third vaccination leading to a memory B cell pool targeting not only SARS-CoV-2 variants but other sarbecoviruses remains to be determined.

In summary, our study demonstrates potent, broad, and durable immunity elicited by the AS03-adjuvanted RBD-I53-50 vaccine platform, which confers protection against Omicron at least until 6 months after vaccination. These results have important implications for the vaccine currently in phase 3 clinical trials.

## Acknowledgements

This study was supported by the Bill and Melinda Gates Foundation INV– 018675; OPP1156262 to D.V. and N.P.K.; National Institute of Allergy and Infectious Diseases (DP1AI158186 and HHSN272201700059C to D.V.), a Pew Biomedical Scholars Award (D.V.), an Investigators in the Pathogenesis of Infectious Disease Awards from the Burroughs Wellcome Fund (D.V.), Fast Grants (D.V.); a generous gift from the Audacious Project (N.P.K.), a generous gift from J. Green and M. Halperin, a generous gift from the Hanauer family, the Defense Threat Reduction Agency (HDTRA1-18-1-0001 to N.P.K., Fast grants (T.T.W.), National Institutes of Health U19 AI111825 (T.T.W.), U54 CA260517 (T.T.W.). D.V. is an Investigator of the Howard Hughes Medical Institute. This project has been funded in whole or in part with Federal funds from the National Institute of Allergy and Infectious Diseases, National Institutes of Health, Department of Health, and Human Services, under Contract No. 75N93021C00016 and 75N93019C00065 to A.S. We thank the staff at the New Iberia Research Center, UL Lafayette, and the Tulane National Primate Research Center for conducting the animal studies; Robert Seder and Daniel Douek for the N gene probe; and L. Stuart at the Bill and Melinda Gates Foundation for inputs and insights throughout the study; all the members of GSK, Dynavax and Seppic for critical reading of the manuscript. We acknowledge the generous help of Rashmi Ravichandran for protein production. The schematic was made using BioRender.

## Author contributions

B.P., H.K. and P.S.A. conceptualized the study; B.P., and P.S.A. designed the study and were responsible for overall conduct of the study; P.S.A., Y.F., and M.H., performed binding antibody assays and T cell assays.; Y.F. performed memory B cell staining; U.A. and V.V.E. performed pseudovirus and live-virus neutralization assays; V.I.Z. calculated antibody half-lives; P.P.A., N. G., K.W.M. G, B. M. T., N.J.N., B.J.B., R.P.B., J.D., and K.R. performed challenge and post-challenge experiments; M.C.M. prepared antigen; K.R., L.S.S., D.E.F., N.R.D.A. processed and purified pre-challenge PBMC, plasma and sera samples; A.G. and A.S. provided peptides for T cell assays; J.F., and F.V., organized the NHP study; D.T.O., R.V.D.M., and R.R. provided AS03 and inputs to the study; F.V., J.R., M.S.S., D.V., T.W., N.P.K. and B.P. supervised the experiments; P.S.A., and B.P. were responsible for the formal analysis of all datasets and preparation of figures; P.S.A., and B.P. wrote the manuscript with suggestions and assistance from all co-authors. All the authors read and accepted the final contents of the manuscript.

## Declaration of interests

B.P. serves on the External Immunology Board of GlaxoSmithKline, and on the Scientific Advisory Board of Sanofi, Medicago, CircBio and Boehringer-Ingelheim. D.T.O., R.V.D.M., and R.R. are employees of GSK group of companies. H.K. is an employee of Bill and Melinda Gates Foundation. C.-A.S. is a consultant for Gritstone Bio, Flow Pharma, Arcturus Therapeutics, ImmunoScape, CellCarta, Avalia, Moderna, Fortress and Repertoire. LJI has filed for patent protection for various aspects of T cell epitope and vaccine design work.

## Methods

### Animal subjects and experimentation

Twenty one male rhesus macaques (*Macaca mulatta*) of Indian origin, aged 5 – 15 years, including 11 animals from the previous study(Arunachalam et al., 2021) were housed and maintained as per National Institutes of Health (NIH) guidelines at the New Iberia Research Center (NIRC) of the University of Louisiana at Lafayette in accordance with the rules and regulations of the Committee on the Care and Use of Laboratory Animal Resources. The entire study (protocol 2020-8808-015) was reviewed and approved by the University of Louisiana at Lafayette Institutional Animal Care and Use Committee (IACUC). All animals were negative for simian immunodeficiency virus, simian T cell leukaemia virus and simian retrovirus. For the challenge, the animals were transferred to the Regional Biosafety Level 3 facility at the Tulane National Primate Research Center, where the study was reviewed and approved by the Tulane University IACUC (protocol 3930).

### RBD-16GS-I53-50 nanoparticle immunogen production

Nanoparticle immunogen components and nanoparticles were produced as previously described in detail(Walls et al., 2020) with the exception that the RBD-Wu was in a buffer containing 50 mM Tris pH 8, 150 mM NaCl, 100 mM L-arginine and 5% w/v sucrose and the RBD-β was in a buffer containing 50 mM Tris pH 8, 150 mM NaCl, 100 mM L-arginine and 5% v/v glycerol.

### Adjuvant formulation and immunization

AS03 is an oil-in-water emulsion that contains 11.86 mg α-tocopherol, 10.69 mg squalene, and 4.86 mg polysorbate 80 (Tween-80) in PBS. For each dose, the nanoparticle immunogens were diluted to 50 μg ml^−1^ (SARS-CoV-2 antigen component) in 250 μl of Tris-buffered saline (TBS) and mixed with an equal volume of AS03. The dose of AS03 was 50% v/v (equivalent of one human dose) All immunizations were administered via the intramuscular route in right forelimbs. The volume of each dose was 0.5 ml.

### Anti-Spike electrochemiluminescence (ECL) binding ELISA

Anti-Spike IgG titers were measured using V-plex COVID-19 panel 23 from Mesoscale Discovery (Cat #K15567U). The assay was performed as per the manufacturer’s instructions. Briefly, the multi-spot 96 well plates were blocked in 0.15 ml of blocking solution with shaking at 700 rpm at room temperature. After 30 min of incubation, 50 μl of sera or BAL samples diluted in antibody diluent solution, and serially diluted calibrator solution was added to each plate in the designated wells and incubated at room temperature for 2 h with shaking. After 2 h of incubation, the plates were washed and 50 μl of Sulfo-tag conjugated anti-IgG was added, and the plates were incubated at room temperature for 1 h. After incubation, the plates were washed and 0.15 ml of MSD-Gold read buffer was added. The plates were immediately read using the MSD instrument. The unknown concentrations were extrapolated using a standard curve drawn using the calibrators in each plate and presented as relative MSD Arbitrary Units (AU)/ml.

### Pseudovirus production and neutralization assay

VSV-based GFP/nanoluciferase-expressing SARS-CoV-2 pseudoviruses were produced as described previously**(Sievers et al., 2022)**. VSV-ΔG-GFP/nanoluciferase and plasmids encoding spike genes of SARS-CoV-2 Wuhan (SΔ19), Beta (B.1.351), and Omicron (B.1.529) were provided by Dr. Gene S. Tan (J. Craig Venter Institute, La Jolla, CA). To perform neutralization assay, Vero E6-TMPRSS2-T2A-ACE2 cells (BEI Resources, NIAID; NR-54970) were seeded at a density of 1×10^4^ per well in half area 96-well black opaque plates (Greiner Bio-One) and were grown overnight at 37°C in a 5% CO2 atmosphere. Serum samples were 5-fold serially diluted using the infection medium (DMEM supplemented with 2% FBS and 100 U/mL Penicillin-Streptomycin) in duplicates. Diluted serum samples were then mixed with an equal volume of Wuhan, Beta, or Omicron pseudoviruses, diluted in infection medium at an amount of 200-400 focus-forming units/mL per well, followed by incubation at 37°C for 1 hour. Subsequently, immune complexes were added onto the monolayers of PBS-washed Vero E6-TMPRSS2-T2A-ACE2 cells and incubated at 37°C. At 18 hours post-incubation, supernatants were removed, cells were washed once with PBS, and nanoluciferase enzymatic activities were measured using the Nano-Glo Luciferase Assay System (Promega) and a SpectraMax iD3 multi-mode microplate reader. Percent inhibition values were calculated by subtracting the percent infection from 100. Non-linear curves and IC_50_ values were determined using GraphPad Prism.

### Viruses and cells

VeroE6-TMPRSS2 cells were described previously and cultured in complete DMEM in the presence of Gibco Puromycin 10 mg/ml (#A11138-03). nCoV/USA_WA1/2020 (WA/1) was propagated from an infectious SARS-CoV-2 clone as previously described(Xie et al., 2020). icSARS-CoV-2 was passaged once to generate a working stock. The hCoV-19/USA/MD-HP01542/2021 (B.1.351) was provided by Dr. Andy Pekosz (Johns Hopkins University, Baltimore, MD) and was propagated in Vero-TMPRSS2 cells(Pegu et al., 2021). hCoV19/EHC_C19_2811C (referred to as the B.1.1.529 variant) was derived from a mid-turbinate nasal swab collected in December 2021. This SARS-CoV-2 genome is available under GISAID accession number EPI_ISL_7171744. All viruses used in the FRNT assay were deep sequenced and confirmed as previously described(Edara et al., 2021).

### Focus Reduction Neutralization Test

FRNT assays were performed as previously described(Edara et al., 2022; Edara et al., 2021; Vanderheiden et al., 2020). Briefly, samples were diluted at 3-fold in 8 serial dilutions using DMEM in duplicates with an initial dilution of 1:10 in a total volume of 60 μl. Serially diluted samples were incubated with an equal volume of WA1, B.1.351, or B.1.1.529 (100-200 foci per well based on the target cell) at 37° C for 45 minutes in a round-bottomed 96-well culture plate. The antibody-virus mixture was then added to VeroE6-TMPRSS2 cells and incubated at 37° C for 1 hour. Post-incubation, the antibody-virus mixture was removed and 100 μl of pre-warmed 0.85% methylcellulose (Sigma-Aldrich, #M0512-250G) overlay was added to each well. Plates were incubated at 37° C for either 18 hours (WA1 or B.1.351) or 40 hours (B.1.1.529) and the methylcellulose overlay was removed and washed six times with PBS. Cells were fixed with 2% paraformaldehyde in PBS for 30 minutes. Following fixation, plates were washed twice with PBS and permeabilization buffer (0.1% BSA and 0.1% Saponin in PBS) was added to permeabilized cells for at least 20 minutes. Cells were incubated with an anti-SARS-CoV spike primary antibody directly conjugated to Alexaflour-647 (CR3022-AF647) for 4 hours at room temperature or overnight at 4° C. Cells were washed three times in PBS and foci were visualized on an ELISPOT reader. Antibody neutralization was quantified by counting the number of foci for each sample using the Viridot program(Katzelnick et al., 2018). The neutralization titers were calculated as follows: 1 - (ratio of the mean number of foci in the presence of sera and foci at the highest dilution of respective sera sample). Each specimen was tested in duplicate. The FRNT-50 titers were interpolated using a 4-parameter nonlinear regression in GraphPad Prism 9.2.0. Samples that do not neutralize at the limit of detection at 50% are plotted at 10 and was used for geometric mean and fold-change calculations.

### Antibody half-life calculations

Mixed-effects models implemented in MonolixSuite 2021R1 (Lixoft) were used to estimate the corresponding half-lives of antigen-specific antibodies. The equations dAb/dt=-*k*\*Ab (for the exponential decay model) and dAb/dt=-*k*/t*Ab (for the power law decay model) were fitted to the longitudinal data starting from day 21 after the second or third vaccine doses, where Ab is the antibody level and *k* is the exponential or power law decay rates, respectively. The corresponding half-lives were calculated as t_1/2_=ln(2)/*k* for the exponential decay model or t_1/2_ (estimated from a given time T) = T(2^1/*k*^ −1) for the power law decay model. For the power law model the half-lives were estimated at day T=100 days after the second or third vaccine doses. Longitudinal data from the comparable three time points were used to estimate the decay rates after the second dose (days 21, 98, 180) and the third dose (days 21, 90, 123-151). The individual-level parameters were lognormally distributed for the initial Ab level (at day 21) and normally distributed for the decay rate *k* with an assumption of no correlations between the random effects. We assumed multiplicative independent lognormal observation error. The estimation of the population parameters was performed using the Stochastic Approximation Expectation-Maximization (SAEM) algorithm.

### Intracellular cytokine staining assay

Antigen-specific T cell responses were measured using the intracellular cytokine staining assay. Live frozen PBMCs were revived, counted and resuspended at a density of 10^6^ live cells per ml in complete RPMI (RPMI supplemented with 10% FBS and antibiotics). The cells were rested overnight at 37 °C in a CO_2_ incubator. Next morning, the cells were counted again, resuspended at a density of 12 10^6^ per ml in complete RPMI and 100 μl of cell suspension containing 1.2 10^6^ cells was added to each well of a 96-well round-bottomed tissue culture plate. Each sample was treated with 3 or 4 conditions depending on cell numbers, no stimulation, a peptide pool spanning the Spike protein of the ancestral Wu strain, Omicron variant and Beta variant (where cell numbers permitted) in the presence of 1 μg ml^−1^ of anti-CD28 (clone CD28.2, BD Biosciences) and anti-CD49d (clone 9F10, BD Biosciences) as well as anti-CXCR3 and anti-CXCR5. The details of peptide synthesis and purity are described previously(Tarke et al., 2022). All samples contained 0.5% v/v DMSO in total volume of 200 μl per well. The samples were incubated at 37°C in CO2 incubators for 2 h before addition of 10 μg ml^−1^ brefeldin A. The cells were incubated for an additional 4 h. The cells were washed with PBS and stained with Zombie UV fixable viability dye (Biolegend). The cells were washed with PBS containing 5% FCS, before the addition of surface antibody. The cells were stained for 20 min at 4 °C in 100 μl volume. Subsequently, the cells were washed, fixed and permeabilized with cytofix/cytoperm buffer (BD Biosciences) for 20 min. The permeabilized cells were stained with intracellular cytokine staining antibodies for 20 min at room temperature in 1× perm/wash buffer (BD Biosciences). Cells were then washed twice with perm/wash buffer and once with staining buffer before acquisition using the BD Symphony Flow Cytometer and the associated BD FACS Diva software. All flow cytometry data were analysed using Flowjo software v10 (TreeStar Inc.).

### Spike-specific memory B cell staining

Cryopreserved PBMCs were thawed and washed twice with 10 ml of FACS buffer (1 x PBS containing 2% FBS and 1 mM EDTA) and resuspended in 100 μl of 1x PBS containing Zombie UV live/dead dye at 1:200 dilution (BioLegend, 423108) and incubate at room temperature for 15 minutes. Following washing, cells were incubated with an antibody cocktail for 1 hour protected from light on ice. The following antibodies were used: IgD PE (Southern Biotech, 2030-09), IgM PerCP-Cy5.5 (BioLegend, 314512), CD20 APC-H7 (BD Biosciences, 560734), CD27 PE-Cy7 (BioLegend, 302838), CD14 BV650 (BioLegend, 301836), CD16 BV650 (BioLegend, 302042), IgG BUV496 (BD Biosciences, 741172), CD3 BV650 (BD Biosciences, 563916), CD21 PE-CF594 (BD Biosciences, 563474) and Alexa Fluor 488-labeled Beta spike (antibodies-online, ABIN6963740), Alexa Fluor 647 labeled Omicron spike (SinoBiological, 40589-V08H26) and BV421 labeled Wuhan Spike (SinoBiological, 40589-V27B-B). All antibodies were used as the manufacturer’s instruction and the final concentration of each probe was 0.1 ug/ml. Cells were washed twice in FACS buffer and immediately acquired on a BD FACS Aria III and the Flowjo was used for data analysis.

### Viral challenge

The details of the Omicron challenge stock and the sequencing confirmation have been described previously(Gagne et al., 2022). We used the same challenge stock. Animals were inoculated via the intratracheal and intranasal routes with a total of 2 × 10^6^ PFUs of SARS-CoV-2 Omicron. The inoculum was divided into two equal parts, and 1 × 10^6^ PFUs was inoculated intra-tracheally and 0.5 × 10^6^ PFUs was inoculated into each nostril in 0.5 ml volume.

### Sampling of nares

The monkeys were anaesthetized and placed in dorsal recumbency or a chair designed to maintain an upright posture. Sterile swabs were gently inserted into the nares. Once inserted, the sponge or swab was rotated several times within the cavity or region and immediately withdrawn.

### BAL collection and processing

The animals were anaesthetized using Telazol and placed in a chair designed specifically for the proper positioning for BAL procedures. A local anaesthetic (2% lidocaine) may be applied to the larynx at the discretion of the veterinarian. A laryngoscope was used to visualize the epiglottis and larynx. A feeding tube was carefully introduced into the trachea after which the stylet was removed. The tube was advanced further into the trachea until slight resistance was encountered. The tube was slightly retracted and the syringe attached. Aliquots of warmed normal saline were instilled into the bronchus. The saline was aspirated between each lavage before a new aliquot was instilled. When the procedure was complete, the monkey was placed in right lateral recumbency. The monkey was carefully monitored, with observation of the heart rate, respiratory rate and effort, and mucous membrane colour. An oxygen facemask may be used following the procedure at the discretion of the veterinarian. The monkey was returned to its cage, positioned on the cage floor in right lateral recumbency and was monitored closely until recovery is complete. The BAL samples were filtered twice via 100-μl strainers and collected in 50-ml centrifuge tubes.

The samples were centrifuged at 300*g* for 10 min at 4 °C. The supernatant was transferred into new tubes, aliquoted and stored at −80 °C until downstream processing.

### Viral load

Quantitative RT–qPCR for the subgenomic (sg) RNA encoding the nucleocapsid (N) protein was performed using the primers, probes and conditions described recently(Gagne et al., 2022). The envelope (E) gene subgenomic PCR was performed as described previously(Arunachalam et al., 2021; Wolfel et al., 2020). Primers and probes for the N subgenomic qRT–PCR were as follows: forward 5’-CGATCTCTTGTAGATCTGTTCTC-3’, reverse 5’-GGTGAACCAAGACGCAGTAT-3’, probe 5’-FAM-CGATCAAAACAACGTCGGCCCC-BHQ1-3’. Both PCRs were run in a 20 μl volume containing 5 μl sample, 900 nM primers, 250 nM probe with TaqPath 1-step RT-qPCR master mix, CG (Thermo Fisher Scientific). The PCR conditions were 2 min at 25 °C for uracil *N*-glycosylase incubation, 15 min at 50 °C for reverse transcription, 2 min at 95 °C (Taq activation), followed by 40 cycles of 95 °C for 3 s (denaturation) and 60 °C for 30 s (annealing and elongation).

### Statistics and data visualization

The difference between any two groups at a time point was measured using a two-tailed nonparametric Mann–Whitney unpaired rank-sum test. The difference between time points within a group was measured using a Wilcoxon matched-pairs signed-rank test. All correlations were Spearman’s correlations based on ranks. All the statistical analyses were performed using GraphPad Prism v.9.0.0 or R version 3.6.1. All the figures were made in GraphPad Prism or R and organized in Adobe Illustrator.

### Reporting summary

Further information on research design is available in the Nature Research Reporting Summary linked to this paper.

## Data availability

All data from the study are included in the manuscript and associated files.

## Figure legends

**Supplementary Fig. 1.**
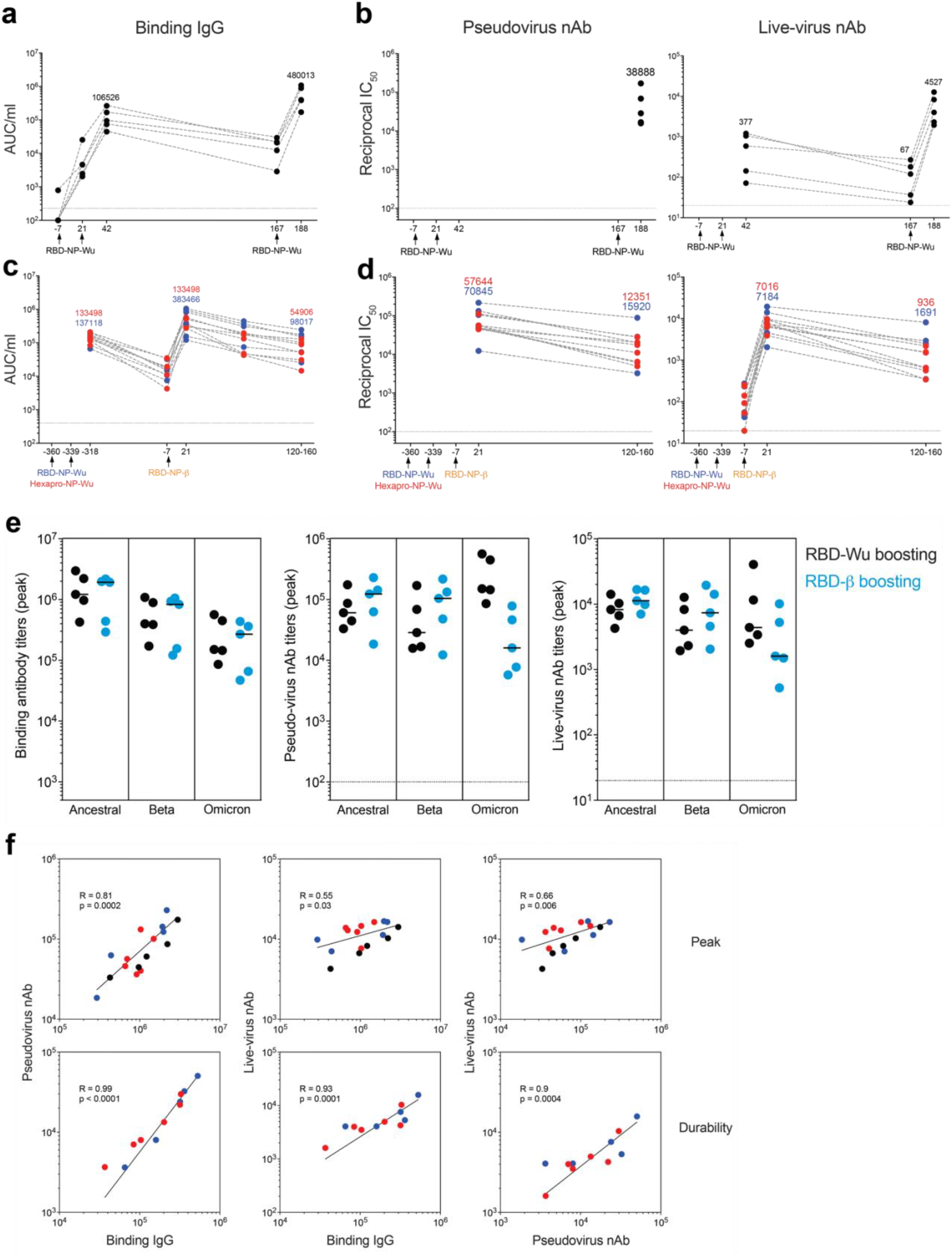
Serum antibody responses induced by AS03-adjuvanted RBD-nanoparticle vaccination. **a - b**, Binding IgG (**a**) and neutralizing antibody (**b**) responses against the Beta variant in the RBD-Wu/RBD-Wu/RBD-Wu group. **c - d**, Binding IgG (**c**) and neutralizing antibody (**d**) responses against the Beta variant in the RBD-Wu/RBD-Wu/RBD-β (blue) and HexaPro/HexaPro/RBD-β groups (red). The numbers indicate geometric mean titers. **e**, Antibody titers in the animals immunized with RBD-Wu/RBD-Wu/RBD-Wu (black, n = 5) or RBD-Wu/RBD-Wu/RBD-β (blue, n = 5) groups. **f**, Spearman’s correlations between Spike-binding IgG, pseudovirus nAb and live-virus nAb titers.

**Supplementary Fig. 2.**
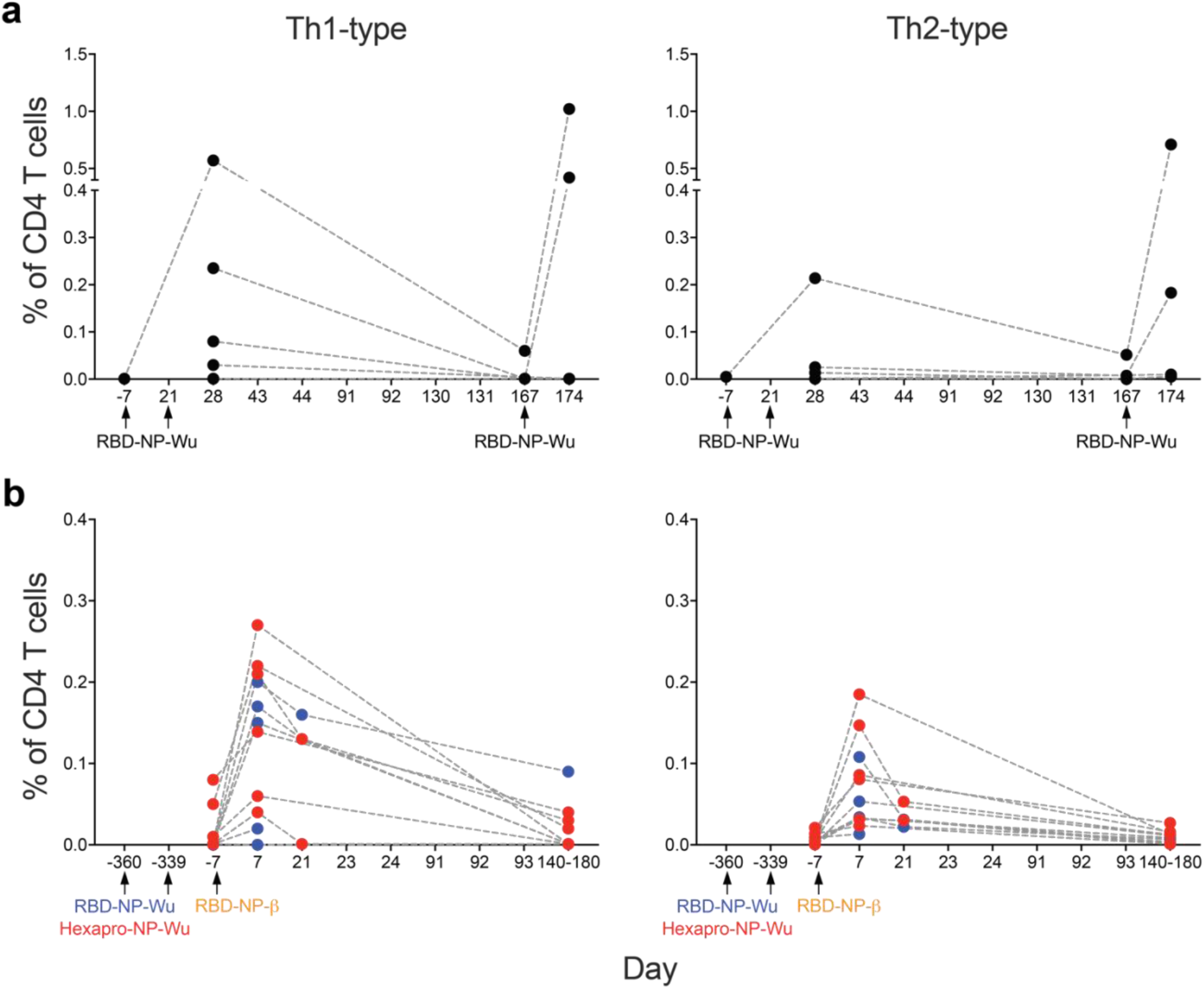
T cell responses induced by AS03-adjuvanted RBD-nanoparticle vaccination. **a - b**, Frequency of Spike-specific CD4 T cell responses against ancestral (left panel) and Omicron (right panel) strains in the RBD-Wu/RBD-Wu/RBD-Wu group (**a**) and the RBD-Wu/RBD-Wu/RBD-β (blue) and HexaPro/HexaPro/RBD-β (red) groups (**b**). CD4 T cells secreting IL-2, IFN-γ, or TNF are plotted as Th1-type responses and IL-4-producing CD4 T cells are shown as Th2-type responses.

**Supplementary Fig. 3.**
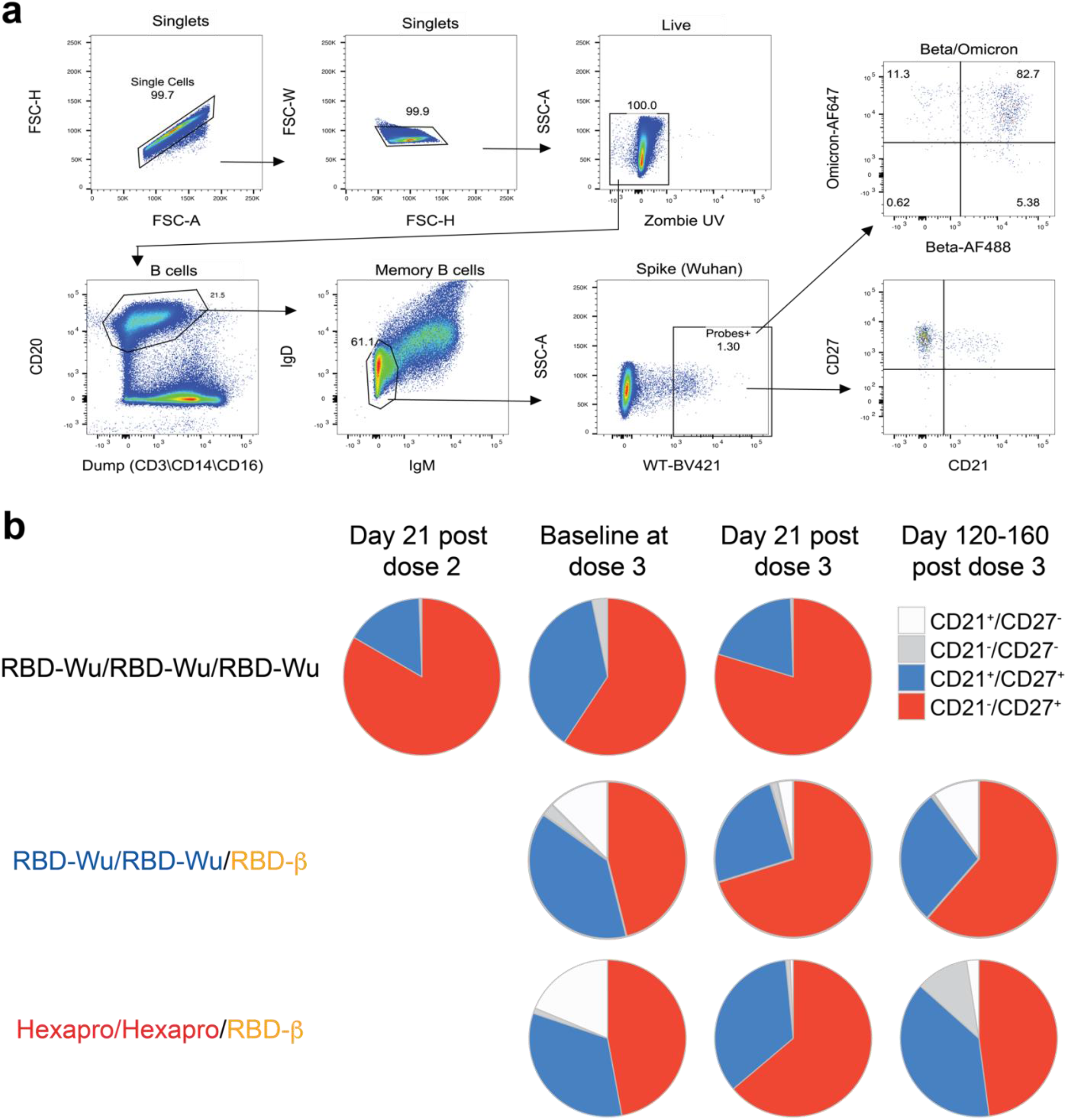
Memory B cell responses induced by AS03-adjuvanted RBD-nanoparticle vaccination. **a**, Flow cytometry gating strategy to identify Spike-specific memory B cells. **b**, Pie charts showing proportion of Spike-specific memory B cells expressing CD21 and CD27 as indicated in the legend.

